# Heterozygous Cell Models of STAT1 Gain-of-Function Reveal a Broad Spectrum of Interferon-Signature Gene Transcriptional Responses

**DOI:** 10.1101/2020.11.09.375097

**Authors:** Ori Scott, Kyle Lindsay, Steven Erwood, Chaim M. Roifman, Ronald D. Cohn, Evgueni A. Ivakine

## Abstract

Signal Transducer and Activator of Transcription 1 (STAT1) gain-of-function (GOF) is an autosomal dominant immune disorder marked by wide infectious predisposition, autoimmunity, vascular disease and malignancy. Its molecular hallmark, elevated phospho-STAT1 (pSTAT1) following interferon (IFN) stimulation, is seen consistently in all patients and may not fully account for the broad phenotypic spectrum associated with this disorder. While over 100 mutations have been implicated in STAT1 GOF, genotype-phenotype correlation remains limited, and current overexpression models may be of limited use in gene expression studies. We generated heterozygous mutants in diploid HAP1 cells using CRISPR/Cas9 base-editing, targeting the endogenous *STAT1* gene. Our models recapitulated the molecular phenotype of elevated pSTAT1, and were used to characterize the expression of five IFN-stimulated genes under a number of conditions. At baseline, transcriptional polarization was evident among mutants compared with wild type, and this was maintained following prolonged serum starvation. This suggests a possible role for unphosphorylated STAT1 in the pathogenesis of STAT1 GOF. Following stimulation with IFN*α* or IFNγ, differential patterns of gene expression emerged among mutants, including both gain and loss of transcriptional function. This work highlights the importance of modelling heterozygous conditions, and in particular transcription factor-related disorders, in a manner which accurately reflects patient genotype and molecular signature. Furthermore, we propose a complex and multifactorial transcriptional profile associated with various *STAT1* mutations, adding to global efforts in establishing STAT1 GOF genotype-phenotype correlation and enhancing our understanding of disease pathogenesis.

## INTRODUCTION

Signal Transducer and Activator of Transcription (STAT) is a family of 7 structurally homologous transcription factors, activated downstream of various cytokine, growth factor and hormone receptors. At rest, STAT molecules are found in a latent state in the cytoplasm. After receptor ligation, canonical STAT activation follows a common sequence, starting with recruitment of tyrosine kinases from the Janus Kinase (JAK) family, which phosphorylate the cytoplasmic portion of the receptor to form a docking site for STAT. This is followed by STAT recruitment, tyrosine phosphorylation and multimerization to form active transcription factors which then migrate to the nucleus.^1–5^ Within the STAT family, STAT1 is pivotal in mediating transcriptional responses to cytokines of the interferon (IFN) family, as well as interleukin-27 (IL-27). This is achieved by the formation of transcription complexes, known as interferon-stimulated gene factor 3 (ISGF3) and gamma activating factor (GAF). ISGF3 is a hetero-trimer consisting of STAT1, STAT2 and IFN-regulatory factor 9 (IRF9). It is primarily formed in the context of type I and III IFN stimulation, and binds to interferon-stimulated response element (ISRE) to regulate gene expression. In contrast, GAF is a STAT1 homo-dimer, predominantly activated in response to type II IFN and IL-27, which exerts its transcriptional activity by binding to gamma-activating sequence (GAS) within gene promoters.^2–6^

Monogenic defects in the *STAT1* gene have been implicated in three distinct human disorders to date. Autosomal recessive complete loss of function (LOF) leads to severe and early-onset susceptibility to viral and Mycobacterial infections. Individuals harbouring two hypomorphic alleles display a milder form of this disease. ^7–9^ A second entity, caused by heterozygous dominant negative mutations, is characterized by Mendelian Susceptibility to Mycobacterial Disease (MSMD).^10–12^ The third disorder, STAT1 gain-of-function (GOF), was first described among a subset of individuals with chronic mucocutaneous Candidiasis and autoimmune thyroid disease, harboring heterozygous point mutations in *STAT1*.^13,14^ The molecular hallmark of the disease was defined as increased levels of phosphorylated STAT1 (with respect to the Tyrosine 701 residue) in response to IFN stimulation.^14^ Since its first description in 2011, STAT1 GOF has been diagnosed in hundreds of patients, and its phenotypic spectrum expanded.^15,16^ Infectious predisposition includes fungal, bacterial, viral, opportunistic and Mycobacterial infections. Over one third of patients display autoimmune features, with hypothyroidism, type 1 diabetes, and cytopenias being common manifestations. Vascular abnormalities, notably intra-cerebral aneurysms, have been described at an increased frequency compared with the general population. Malignancies, in particular squamous cell carcinoma, are seen in up to 5% of patients.^15–20^

In the decade since STAT1 GOF was first described, strides have been made in characterizing the disorder and its underlying pathophysiology. A prominent example is the impaired Th17 response observed in most patients, which has been linked to predisposition to fungal and bacterial infections.^14,20,21^ From an autoimmune standpoint, impaired type I IFN response has been proposed as a possible contributory mechanism, given the heightened IFN signature associated with other autoimmune and inflammatory conditions.^22–24^ Indeed, alterations in IFN-related gene expression have been found in some patients with STAT1 GOF and clinical features of autoimmunity.^25^ However, many underlying disease mechanisms have remained elusive, and the genotype-phenotype correlation among patients remains poorly defined. A recent review reported that the presence of a severe complication, defined as invasive infection, cancer, symptomatic aneurysm, and in some cases severe autoimmunity, substantially worsens the prognosis and lowers survival.^26^ Unfortunately, our ability to predict which patients would be more prone to developing such complications based on their specific mutations is greatly limited. Therefore, the need for developing tools to study the variability across *STAT1* GOF mutations is dire.

In studying the differences across *STAT1* GOF mutations, the use of cell models offers a well-controlled, accessible and non-invasive tool. Previous studies utilizing over-expression models generated in *STAT1*-null U3 fibrosarcoma cells (and more recently, HEK293 cells), have been instrumental in elucidating differences among mutations with respect to STAT1 phosphorylation kinetics, nuclear migration and accumulation.^14,27–31^ However, such models involve the expression of *STAT1* under an exogenous promoter, and do not capture the heterozygous nature of the mutation. In the context of a delicately-regulated transcription factor, over-expression models may therefore be limited in their portrayal of gene expression patterns downstream of STAT1. The current study describes our use of the Clustered Regularly Interspaced Short Palindromic Repeats (CRISPR)/Cas9 system, and in particular CRISPR/Cas9 base-editing, to generate a series of heterozygous cell models harbouring known GOF point mutations, within the endogenous *STAT1* gene. We further use these models to show that *STAT1* GOF mutations result in different patterns of interferon-stimulated gene (ISG) expression, both at baseline and following stimulation with IFN*α* or IFNγ. We propose that such models may enhance our understanding of this intricate immune disorder, as well as genotype-phenotype correlation among various *STAT1* GOF mutations.

## RESULTS

### Diploid HAP1 cells were chosen for heterozygous mutation modelling

For the purpose of our model generation we have used HAP1, a cell line originally derived from the KBM-7 chronic myelogenous leukemia cell line. HAP1 were previously used to study cellular responses to IFN types I, II and III, and have well-characterized transcriptional responses to IFN type I and II.^32–34^ In addition, they are readily amenable to transfection and CRISPR/Cas genome-editing.^35^ Although HAP1 cells are originally near-haploid, like other haploid cell lines they are known to spontaneously diploidize in cell culture over time.^36^ HAP1 cells used in this study underwent cell cycle analysis, comparing their DNA content with that of known diploid cells [wildtype (WT) human fibroblasts]. The mean fluorescence intensity (MFI) ratios of HAP1/fibroblasts for the G_0_/G_1_ and G_2_/M peaks were calculated to be 1.07 and 1.11, respectively, confirming our HAP1 cells to be fully diploid, and therefore suitable for heterozygous mutation modelling (**Supplementary Figure 1**).

### Heterozygous *STAT1* mutants were generated using CRISPR/Cas9 base-editing

The following validated GOF transition mutations were chosen for modelling: E235G,^37^ K278E^38^ (both in the coiled-coil domain), P329L,^39,40^ T385M^16,41–49^ (in the DNA-binding domain), and D517G^15^ (in the linker domain). The dominant negative mutation Y701C,^50^ affecting the JAK-phosphorylated residue Y701, was chosen for comparative modelling as well. A visual representation of modelled mutations within their respective protein domains, as well as the workflow for generating and verifying the *STAT1* mutants, is presented in **Figure 1**. For information regarding clinical manifestations described for each mutation, please refer to **Table I.**

**Figure 1.**
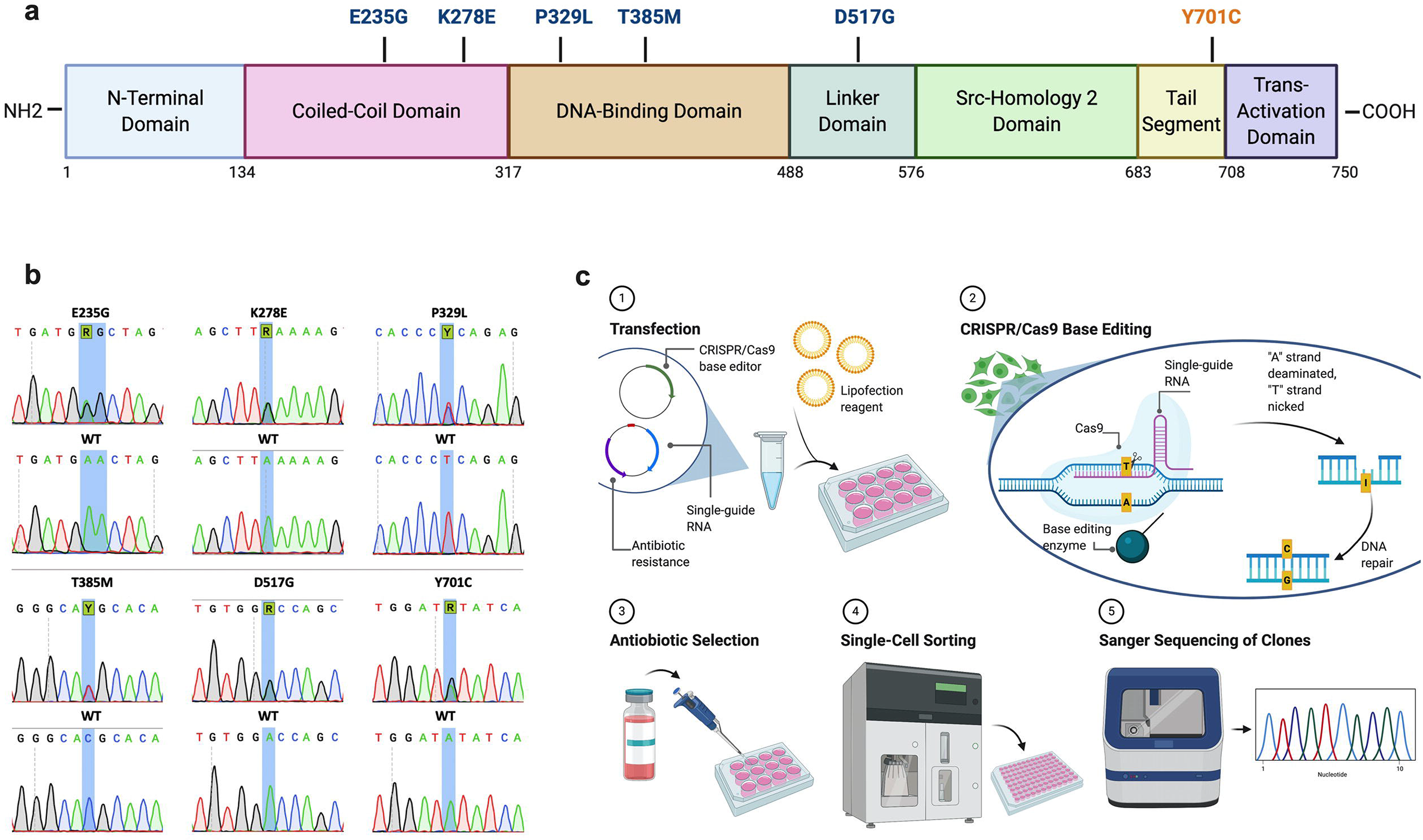
*STAT1* mutant generation. | (a) Schematic representation of *STAT1* mutations selected for modelling and their respectively-affected protein domains. Mutations designated in blue (E235G, K278E, O329L, T385M, D517G) are “gain-of-function” mutations, while orange (Y701C) designates a “loss-of-function” mutation. (b) Overview of the process of heterozygous cell model generation using CRISPR/Cas9 base editing. (c) Sanger sequencing confirmation of generated cell models, compared with their respective wildtype counterparts. Highlighted nucleotides denote the edited base. Note that for E235G, editing of an adjacent “A” on both alleles took place, resulting in a silent bystander mutation. This yielded the trinucleotide change: GAA/GAA→ GAG/GGG, leading to the amino acid substitution: Glutamine/Glutamine→ Glutamine/Glycine. This corresponds with the same amino acid change found in patients affected by the E235G mutation.

**Table I:** Clinical manifestations of *STAT1* mutations modelled in this work.

| **Mutation** | **Designation** | **Fungal infections** | **Bacterial Infections** | **Viral Infections** | **Mycobacterial infections** | **Autoimmunity / Endocrinopathy** | **Vascular abnormalities** | **Malignancy** | **Other** |
| --- | --- | --- | --- | --- | --- | --- | --- | --- | --- |
| E235G (c.704A>G)<sup>37</sup> | AD GOF | CMCC | Sinopulmonary infections, bacterial skin infections with abscess formation | Recurrent HSV labialis | Mycobacterial pneumonia | Alopecia | Carotid and celiac/ splenic artery dissection | HPV+ squamous cell carcinoma, basal cell carcinoma | N.D. |
| K278E (c.832A>G)<sup>38</sup> | AD GOF | CMCC | Bacterial skin infections | Recurrent HSV stomatitis, recurrent Herpes zoster | N.D. | Positive auto-antibodies (ANA, microsomal, anti-TSH, anti-IL-17F) | N.D. | N.D. | N.D. |
| P329L (c.986T>C)<sup>39,40</sup> | AD GOF | CMCC | Yes, no details provided | Recurrent HSV stomatitis | N.D. | Autoimmune hemolytic anemia, red blood cell aplasia | N.D. | N.D. | N.D. |
| T385M (c.1154C>T)<sup>16,41-49</sup> | AD GOF | CMCC, invasive fungal infections | Sinopulmonary infections | EBV, CMV, recurrent severe VZV | Variety of Mycobacterial infections reported | Enteropathy, T1DM, thyroiditis, autoimmune cytopenias, vitiligo, growth hormone deficiency, autoimmune hepatitis | Intracranial aneurysms | N.D. | Recurrent fractures, eczema, chronic adenopathy, multifocal leuko-encephalopathy |
| D517G (c.1550A>G)<sup>15</sup> | AD GOF | CMCC, invasive fungal infections | N.D. | N.D. | N.D. | N.D. | N.D. | N.D. | N.D. |
| Y701C (c.2102A>G)<sup>50</sup> | AD LOF | N.D. | N.D. | N.D. | Invasive Mycobacterial infections | N.D. | N.D. | N.D. | Multifocal osteomyelitis |

AD: Autosomal dominant; ANA: anti-nuclear antibody; CMCC: chronic mucocutaneous Candidiasis; CMV: Cytomegalovirus; EBV: Epstein-Barr virus; GOF: gain-of-function; HPV: Human papillomavirus; HSV: Human Herpesvirus; LOF: loss-of-function; N.D.: not described; T1DM: Type 1 diabetes mellitus; TSH: thyroid stimulating hormone; VZV: Varicella zoster virus

To generate chosen mutations, single-guide RNA (sgRNA) targeting the Cas9 base-editor to the region of interest were cloned into the BPK1520_puroR plasmid. Resultant plasmids, coding for the desired sgRNA as well as a puromycin resistance cassette, were delivered by lipofection into HAP1 cells, concurrently with an additional plasmid coding the respective Cas9 base-editor (SpCas9 ABEmax, SpG CBE4max, or SpRY ABEmax). Following puromycin selection to enrich for transfected cells, base-editing efficiency was evaluated in the bulk-population by sequencing. Overall, editing efficiency ranged from 21 to 77% for the desired target nucleotide. As base editors each have a characteristic “editing window”, adjacent nucleotides to the nucleotide of interest may be prone to “bystander” editing. As an example, if two adjacent adenine residues are both within the editing window for an adenine base editor, both may be targeted and converted to guanine, albeit at different frequencies depending on their position within the editing window. In this study, bystander mutations involving editing of adjacent bases within the editing window occurred at a frequency of 1-38%. Cells from the bulk population were single-cell sorted, and resultant single clones were screened by Sanger sequencing for presence of the desired mutation in a heterozygous state, and for absence of non-silent bystander mutations. For all mutations, the number of clones required to be screened to identify a heterozygous mutant without bystander mutations ranged from 12 to 39. For one mutation, E235G, only clones containing a second, silent mutation in an adjacent base could be identified. However, these clones recapitulated the desired amino acid change and were therefore deemed appropriate for further downstream work. Altogether, all chosen amino acid changes could be modelled using base-editing. For each mutation generated, downstream analysis was carried out on a minimum of 2 independent clones. A summary of base-editing performed in this work, including sgRNA and base-editors used, frequency of editing events in the bulk population, and number of clones required to screen to find a heterozygous mutation is provided in **Table II**.

**Table II:**
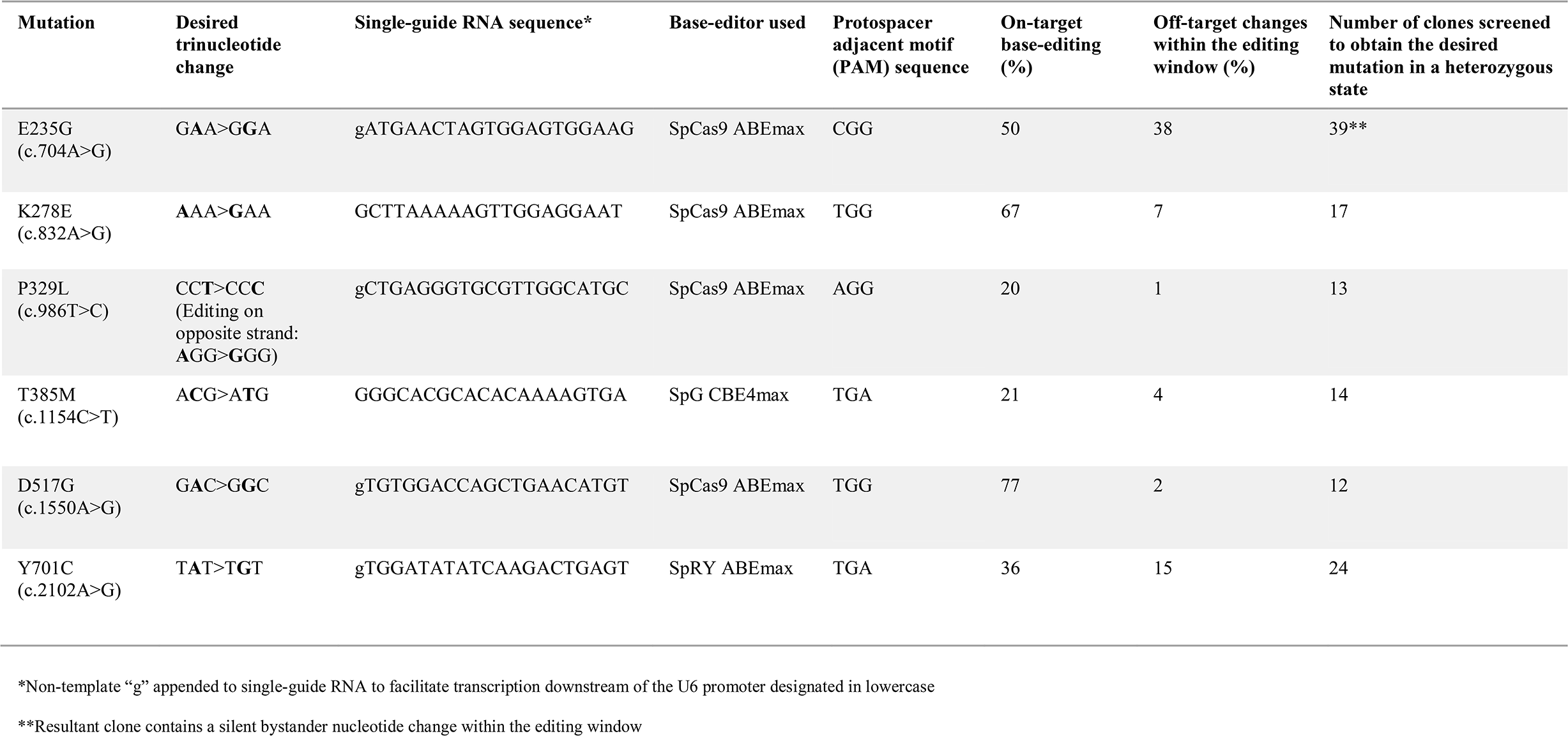
Summary of base-editing features for *STAT1* mutation generation.

### Immunoblotting for pSTAT1 validated the molecular designation of generated *STAT1* mutants

In order to establish the validity of our newly-generated cell models, we proceeded to validate their designation as “GOF” or “LOF” based on Tyrosine-701 phosphorylation in response to IFN. To this end, cells were stimulated with IFNγ at a dose of 10ng/mL for a period of 60 minutes. pSTAT1(Y701) and total STAT1 were measured by immunoblotting at baseline in unstimulated cells, as well as following stimulation. Measurements were repeated over 5 independent experiments and densitometry analysis performed (**Figure 2; Supplementary Figure 2**). At baseline, pSTAT1 was not detectable in any of the samples. Following IFNγ stimulation, levels of pSTAT1 increased across all samples, but were significantly higher across all GOF mutants compared with WT [E235G (*p<0.05*), K278E, P329L, T385M (*p<0.001*), D517G (*p<0.01*); One-way ANOVA with Dunnett’s post-hoc test]. By the same token, pSTAT1 was lower in the Y701C LOF mutant compared with WT after stimulation (*p<0.05*). These results establish that modelled heterozygous *STAT1* mutations in HAP1 cells lead to the same functional consequences with respect to protein phosphorylation as are seen in patients. Total STAT1 was significantly elevated only in the T385M mutant (*p<0.05*), although a trend toward increased total STAT1, not reaching statistical significance, was seen in P329L (*p=0.0635*) and D517G (*p=0.0792*) as well.

**Figure 2.**
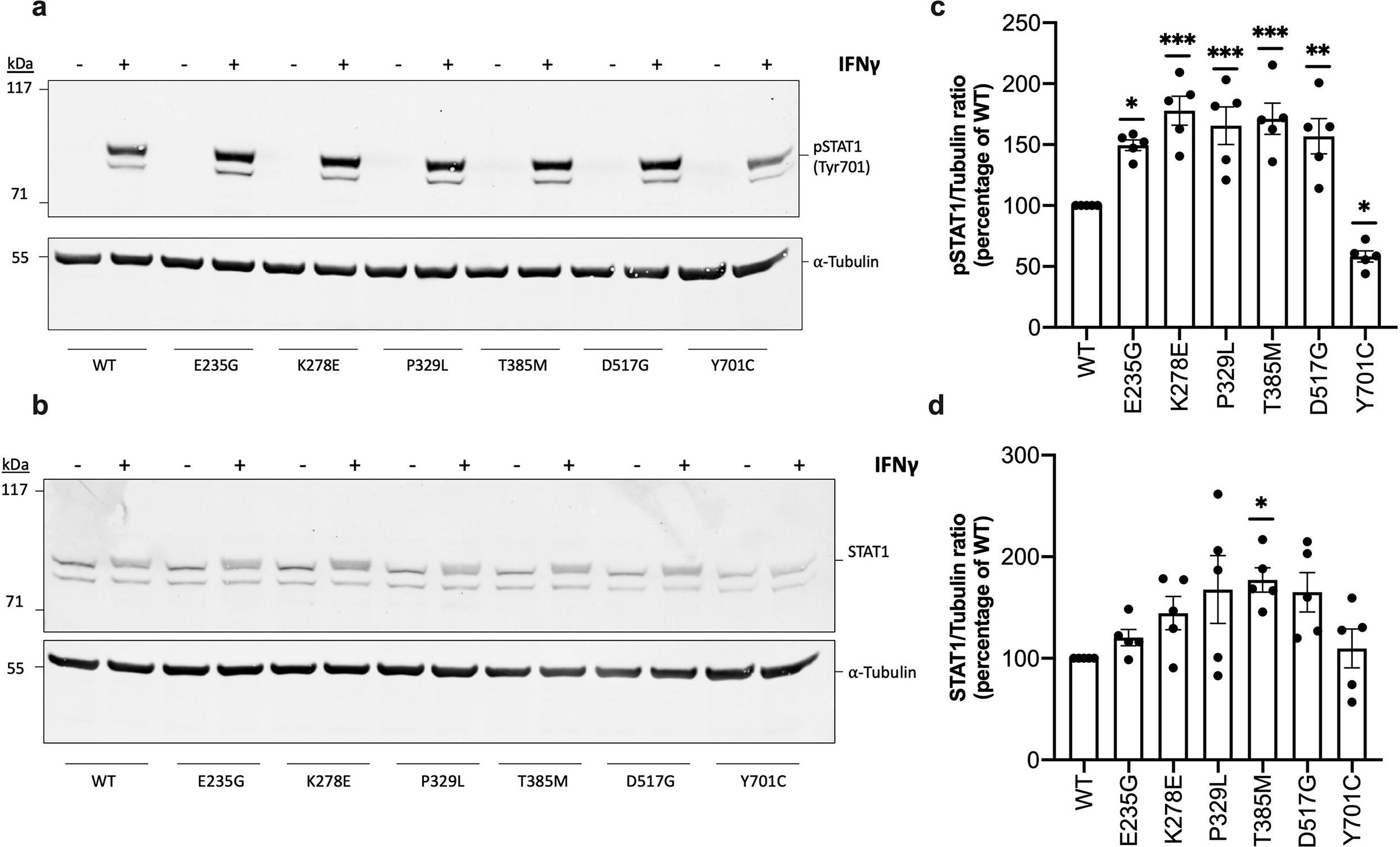
Immunoblot analysis of pSTAT1 (Tyr701) and total STAT1 levels among STAT1 mutants following IFNγ stimulation. | Levels of pSTAT1 (Tyr701; a) and total STAT1 (b) were measured in whole cell protein lysates at baseline and following a 60-minute stimulation with IFNγ (10ng/mL), normalized to a loading control (⍰-Tubulin). Densitometry analysis results from 5 independent experiments were plotted for pSTAT1 (c) and STAT1 levels following IFNγ stimulation (d) and compared with WT. Results are presented as mean + standard error of mean. Statistical analysis: One-way ANOVA with Dunnett’s post-hoc test (* *p<0.05*, ** *p<0.01*, *** *p<0.001*, **** *p<0.0001*).

### Gene expression studies demonstrated baseline polarization among *STAT1* mutants

To evaluate the transcriptional impact of the various *STAT1* mutations, we used quantitative real-time PCR (qRT-PCR) to assessed differences in interferon-stimulated gene (ISG) expression under a number of conditions, including baseline, serum starvation, and stimulation with IFN type I and II. A visual summary of STAT1 signaling associated with the above conditions is presented in **Figure 3**. A set of five ISG was chosen consisting of *GBP1, IFIT2, IRF1, APOL6* and *OAS1*. These genes were selected as they are known to increase in human cells at least 2-fold following either IFN⍰ or IFNγ stimulation.^51^ Moreover, previous studies specifically done in HAP1 cells showed these genes to increase at least 2-fold following stimulation with either type I or II IFN stimulation.^34^

**Figure 3.**
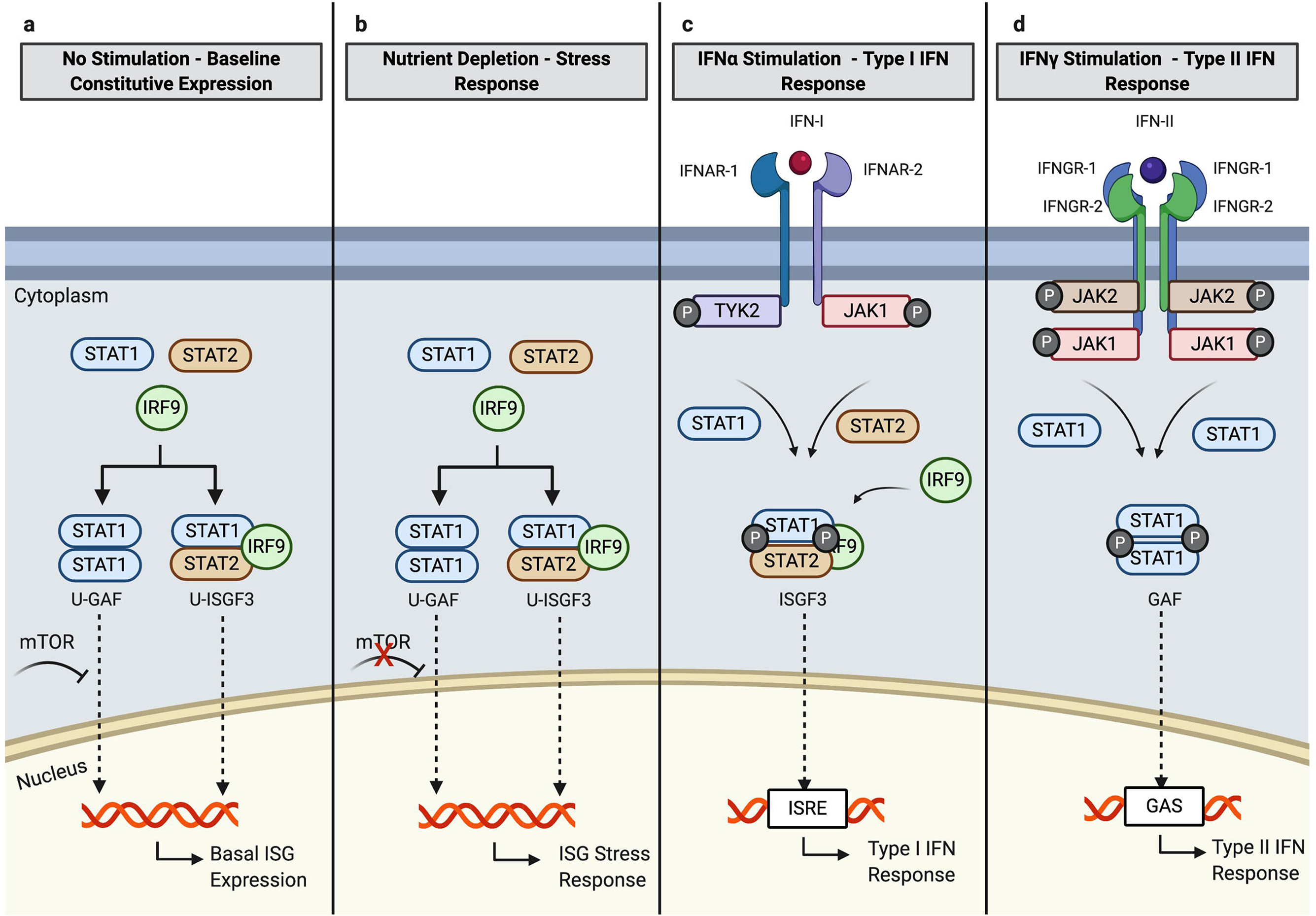
STAT1-associated gene transcription under various experimental conditions. | (a) at baseline, constitutive Interferon-Stimulated Gene (ISG) expression is maintained at a basal level in an interferon (IFN)-independent manner via two unphosphorylated transcription complexes: unphosphorylated gamma activating factor (U-GAF) made up of two STAT1 units, and unphosphorylated interferon-stimulated gene factor 3 (U-ISGF3), made up of STAT1, STAT2 and IRF9. At baseline, mammalian target of rapamycin (mTOR) serves as an inhibitor of STAT1 migration to the nucleus, preventing excessive ISG transcription; (b) following nutrient depletion, mTOR is inhibited, allowing for higher nuclear accumulation unphosphorylated-STAT1 containing transcriptional complexes. This results in an ISG transcriptional response which is IFN-independent; (c) Stimulation with IFN⍰ begins with ligation of the type I IFN receptor, made up of two subunits: IFNAR1 and IFNAR2. Receptor ligation leads to recruitment of the tyrosine kinases TYK2 and JAK1, which cross-phosphorylate each other as well as the intracellular receptor domains. This creates a docking site for STAT1 and STAT2 which bind, are phosphorylated, and along with IRF9 form the transcription complex ISGF3. ISGF3 migrates to the nucleus where it binds to interferon-stimulated response element (ISRE) in gene promoters, resulting in a Type I IFN response; (d) Stimulation with IFNγ begins with ligation of the type II IFN receptor, made up of two subunits each of IFNGR1 and IFNGR2. Receptor ligation leads to recruitment of the tyrosine kinases JAK1 and JAK2, which cross-phosphorylate each other as well as the intracellular receptor domains. This creates a docking site for STAT1 which bind, are phosphorylated, and form the homodimer GAF. GAF migrates to the nucleus where it binds to gamma-activating sequence (GAS) in gene promoters, resulting in a Type II IFN response.

At baseline, significant differences in gene expression among WT and some of the mutants were already noted, involving a mixed pattern of both increased and decreased expression (**Figure 4a, Table III**). The most prominent mutants showing elevated expression (2 genes each) were E235G and P329L; E235G showed increased expression of *GBP1* (*p<0.01*) and *APOL6* (*p<0.001*), while P329L demonstrated increased *APOL6* (*p<0.001*) and *OAS1* expression (*p<0.01*). Other changes noted included reduced *OAS1* expression in Y701C (*p<0.*05), and elevated *IFIT2* expression in D517G (*p<0.01*). No baseline differences were noted between T385M and WT, or between K278E and WT. In order to ensure that the observed transcriptional differences were inherent to the mutants, rather than a result of external cell-culture cytokine/growth factor stimuli, gene expression was measured following 24 hours of serum starvation (**Figure 4b, Table III**). In the context of serum starvation, the mutants E235G and P329L maintained a profile of elevated gene expression. E235G demonstrated increased expression of *GBP1* (*p<0.0001*) and *APOL6* (*p<0.01*) compared to WT, while P329L showed enhanced expression of *APOL6* (*p<0.0001*), *OAS1* (*p<0.001*) and in addition, *IRF1* (*p<0.0001*). Mutants which were previously no different than WT (K278E and T385M) with respect to all genes measured, remained so under serum starvation. In the context of these results, and given that serum starvation in and of itself may impact STAT1 activation and gene transcription in an IFN-independent manner,^52^ we elected to proceed with IFN stimulation experiments under normal cell culture conditions, as described by others.^14,16,25,27,28,30,31^

**Figure 4.**
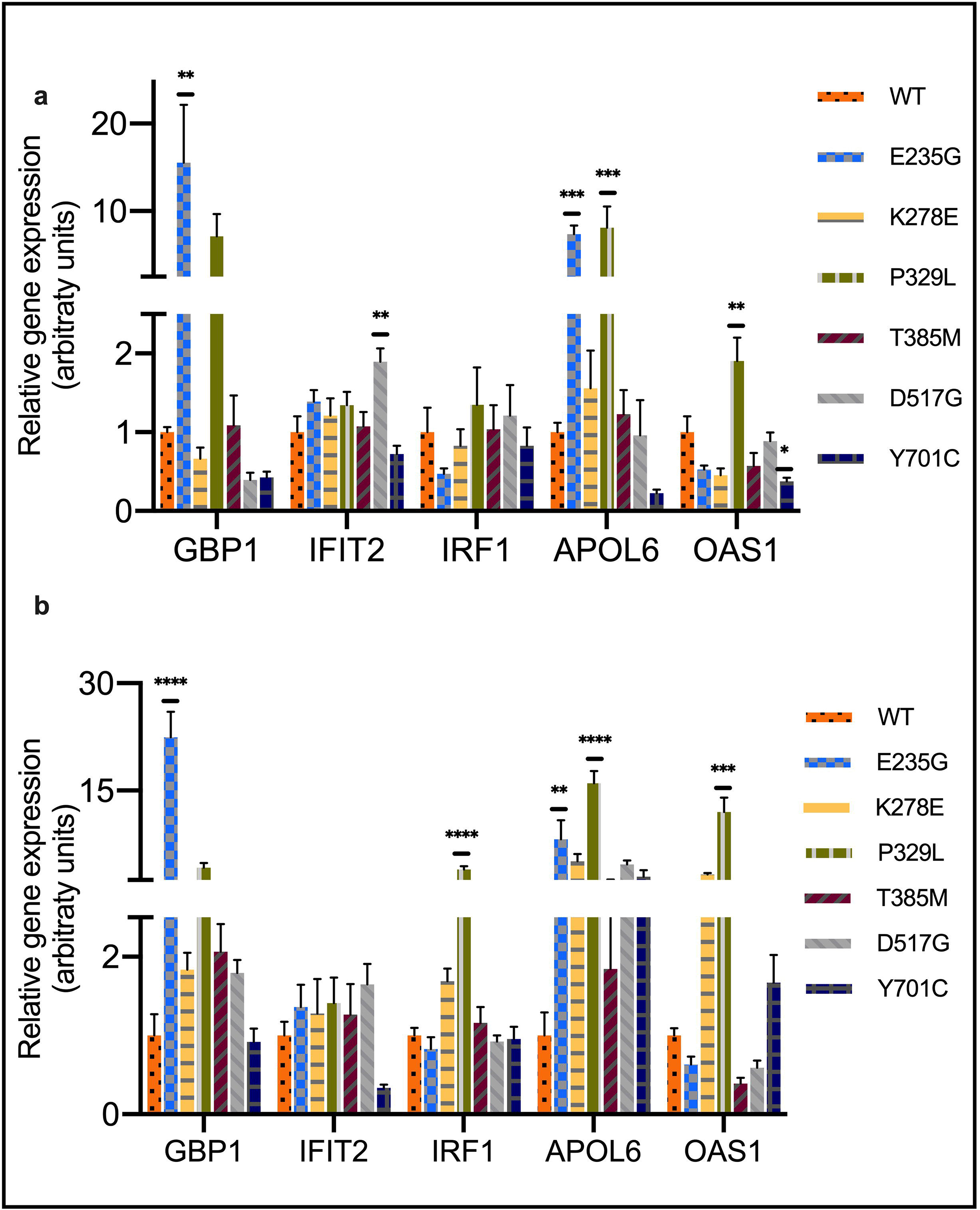
Relative gene expression of Interferon-Stimulated Genes (ISG) at baseline and following serum starvation. **|** (a) mRNA expression levels were measured for five ISG (*GBP1, IFIT2, IRF1, APOL6, OAS1*) at baseline in cells grown under normal serum conditions; (b) mRNA expression levels for five ISG measured in cells following 24-hours of serum starvation. Expression levels were normalized to housekeeping *GAPDH* expression and plotted relatively to wildtype for each experiment. Pooled data from at least 5 experiments are presented. Data are represented as mean ± standard error of mean. Statistical analysis: One-way ANOVA with Dunnett’s post-hoc test (* *p<0.05*, ** *p<0.01*, *** *p<0.001*, **** *p<0.0001*)

**Table III:** Summary of differences in relative expression or fold increase across *STAT1* mutants compared with wildtype under various conditions.

| Mutation | Condition/Stimulation | GBP1 | IFIT2 | IRF1 | APOL6 | OAS1 |
| --- | --- | --- | --- | --- | --- | --- |
| E235G | Baseline (full serum) * | ↑ | NS | NS | ↑ | NS |
|  | Serum starvation ** | ↑ | NS | NS | ↑ | NS |
| | IFNα† | NS | ↑ | ↑ | NS | NS |
| | IFNγ‡ | ↓ | NS | NS | ↓ | ↑ |
| K278E | Baseline (full serum) * | NS | NS | NS | NS | NS |
|  | Serum starvation ** | NS | NS | NS | NS | NS |
| | IFNα† | ↓ | NS | NS | ↓ | ↓ |
| | IFNγ‡ | NS | ↑ | NS | NS | NS |
| P329L | Baseline (full serum) * | NS | NS | NS | ↑ | ↑ |
|  | Serum starvation ** | NS | NS | ↑ | ↑ | ↑ |
| | IFNα† | ↓ | NS | NS | ↓ | ↓ |
| | IFNγ‡ | ↓ | NS | NS | ↓ | NS |
| T385M | Baseline (full serum) * | NS | NS | NS | NS | NS |
|  | Serum starvation ** | NS | NS | NS | NS | NS |
| | IFNα† | ↑ | ↑ | ↑ | ↑ | ↑ |
| | IFNγ‡ | NS | NS | NS | ↓ | NS |
| D517G | Baseline (full serum) * | NS | ↑ | NS | NS | NS |
|  | Serum starvation ** | NS | NS | NS | NS | NS |
| | IFNα† | NS | NS | NS | ↓ | ↓ |
| | IFNγ‡ | NS | NS | NS | ↓ | NS |
| Y701C | Baseline (full serum) * | NS | NS | NS | NS | ↓ |
|  | Serum starvation ** | NS | NS | NS | NS | NS |
| | IFNα† | ↓ | ↓ | ↓ | ↓ | NS |
| | IFNγ‡ | ↓ | ↓ | ↓ | ↓ | NS |

\* Relative gene expression compared with wildtype under full-serum culture conditions without cytokine stimulation
\*\* Relative gene expression compared with wildtype following 24 hours of low-serum conditions without cytokine stimulation
† Fold increase in gene expression from baseline compared with wildtype following 6 hours of stimulation with IFN $\alpha$ (10ng/mL)‡ Fold increase in gene expression from baseline compared with wildtype following 6 hours of stimulation with IFNγ (10ng/mL)
NS: not significant

### *STAT1* mutants displayed a differential response to IFN*α* stimulation involving both loss and gain of transcriptional function

After establishing baseline expression levels, transcriptional responses (fold increase in expression after stimulation) were measured following a 6-hour stimulation with IFN*α* (10ng/mL) (**Figure 5a; Table III**). Of all GOF mutants, only T385M showed an elevated fold change (FC) across all genes measured compared to WT [*GBP1* (*p<0.0001*), *IFIT2* (*p<0.01*), *IRF1* (*p<0.0001*), *APOL6* (*p<0.01*), *OAS1* (*p<0.05*)]. E235G, which had an elevated baseline expression of *GBP1* and *APOL6*, showed an elevated FC in the expression of *IFIT2* (*p<0.0001*) and *IRF1* (*p<0.0001*), with no difference in FC with respect to other genes. In contrast, P329L, which previously showed increased baseline expression of *APOL6* and *OAS1*, showed a reduced FC in the expression of *GBP1* (*p<0.05*), *APOL6* (*p<0.05*) and *OAS1* (*p<0.01*) compared with WT. Decreased FC compared to WT was also seen in K278E [*IRF1* (*p<0.05*), *APOL6* (*p<0.05*), *OAS1* (*p<0.0001*)] and D517G [*APOL6* (*p<0.05*), *OAS1* (*p<0.001*)]. The LOF mutant Y701C was marked by reduced FC across all but 1 gene compared to WT [*GBP1* ([*p<0.01*), *IFIT2* (*p<0.001*), *IRF1* (*p<0.05*), *APOL6* (*p<0.01*)].

**Figure 5.**
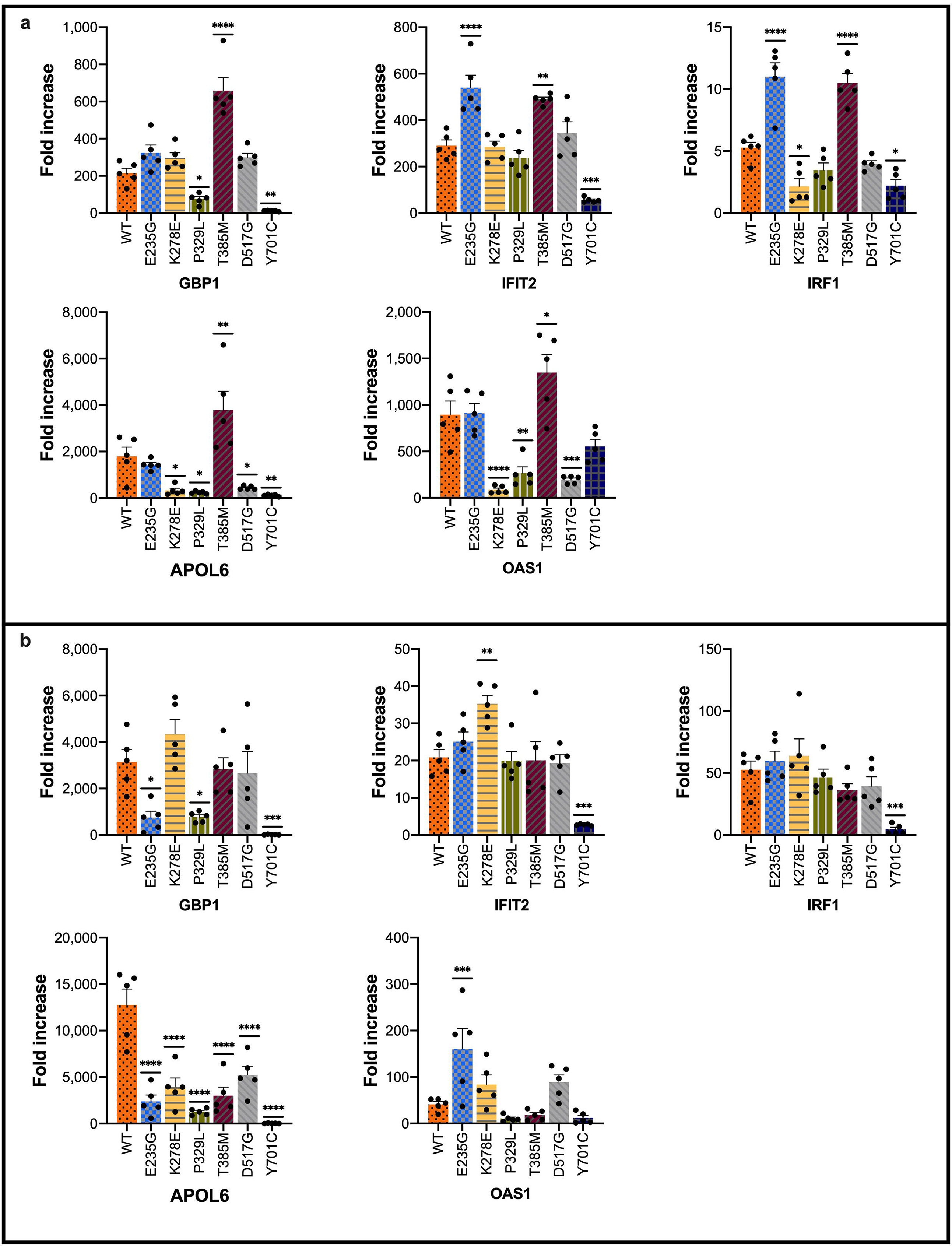
Relative gene expression of Interferon-Stimulated Genes (ISG) following IFN*α* or IFNγ stimulation. **|** mRNA expression levels were measured (*GBP1, IFIT2, IRF1, APOL6, OAS1*) at baseline and following 6 hours of stimulation with IFN⍰ (a) or IFNγ (b). Expression levels were normalized to housekeeping *GAPDH* and plotted as fold increase from baseline. Pooled data from at least 5 experiments are presented. Data are represented as mean ± standard error of mean. Statistical analysis: One-way ANOVA with Dunnett’s post-hoc test (* *p<0.05*, ** *p<0.01*, *** *p<0.001*, **** *p<0.0001*)

### Transcriptional response of *STAT1* mutants to IFNγ stimulation differed from IFN*α* responses

We sought to determine whether transcriptional responses of *STAT1* mutants to IFN*α* could accurately predict their responses to stimulation with IFNγ (**Figure 5b; Table III**). Following a 6-hour stimulation with IFNγ (10ng/mL), 3 mutants showed vastly different transcriptional responses compared with those seen following IFN*α* stimulation. T385M, which previously showed a transcriptional GOF with respect to all genes following IFN⍰ stimulation, now showed a reduced FC of *APOL6* (*p<0.0001*), with no other differences compared with WT. E235G previously demonstrated elevated FC in expression of *IFIT2* and *IRF1* in response to IFN*α*, whereas no differences from WT were seen in these genes with IFNγ stimulation. In contrast, FC of *GBP1* (*p<0.05*) and *APOL6* (*p<0.0001*) were now decreased in E235G, and that of *OAS1* increased (*p<0.001*) compared to WT. One more mutant showing substantial differences in responses to IFNγ and IFN*α* was K278E; while IFN*α* stimulation resulted in reduced FC of *GBP1*, *APOL6* and *OAS1* compared with WT, IFNγ stimulation caused an increased FC in *IFIT2* (*p<0.01*), with no significant differences with respect to other genes. Mutants showing more similar trends in response to both IFNγ and IFN*α* included P329L, D517G, and the LOF mutant Y701C. P329L again showed reduced FC of *GBP1* (*p<0.05*) and *APOL6* (*p<0.0001*), though FC for *OAS1* was no different than WT. D517G demonstrated reduced FC in *APOL6* (*p<0.0001*), but no difference in FC of *OAS1*. Y701C showed consistent responses to IFNγ and IFN*α*, with reduced FC seen again for *GBP1* (*p<0.001*), *IFIT2* (*p<0.001*), *IRF1* (*p<0.001*) and *APOL6* (*p<0.0001*) but no difference in FC of *OAS1*.

## DISCUSSION

The current study demonstrated for the first time the implementation of CRISPR/Cas9 base-editing in creating heterozygous cells models of STAT1 GOF and LOF. Previously, most studies of STAT1 GOF were performed in patient samples, or in overexpression models. Work done in patient-derived samples, be they primary or immortalized cells, has provided a wealth of information regarding pathway alterations associated with STAT1 GOF. However, such samples are a limited resource necessitating access to patients, their obtaining can be invasive, and no perfectly isogenic control is available for comparison. Moreover, as sample collection is typically done after patients have already become symptomatic, it is challenging to exclude variability relating to factors such as concurrent systemic inflammation, infection or immunosuppressive/modulatory treatments. In regards to overexpression models, the majority of studies have employed the *STAT1*-null U3 fibrosarcoma cells (and more recently, HEK293 cells on a WT background).^14,27–31^ These models have been instrumental in studying STAT1 phosphorylation and de-phosphorylation kinetics, as well as migration of STAT1 between the cytoplasm and the nucleus. However, such models are characterized by expression of *STAT1* under an exogenous promoter, and an inaccurate gene dosage. These factors considerably limit the application of overexpression models to the study of precise gene expression and signaling pathway alterations. This limitation is particularly substantial in the case of STAT1, a transcription factor which is under delicate transcriptional control, impacts the expression of other transcription factors, and in itself regulates its own expression.

The current approach of mutant generation via base-editing offers an opportunity to model *STAT1* mutations in a heterozygous manner, and under control of the endogenous gene promoter, resulting in highly relevant cell models for dissecting the molecular pathogenesis of the disease from a transcriptional standpoint. Base-editing is efficient, quick, and enables modelling of a rapidly-expanding repertoire of point mutations.^53–55^ As with most CRISPR/Cas9-based applications, the use of base editing may be limited by the need for a protospacer adjacent motif (PAM) in close proximity to the area of interest. However, with the advent of newly engineered base-editors with extended sequence recognition (such as SpG and SpRY editors used in this work), a wider array of PAMs may now be used in targeting sites for base-editing.^56^ Furthermore, while base-editing was previously limited in its ability to create transversion mutations, recent works have expanded the arsenal of base-editors, now allowing the generation of certain transversions in addition to transitions.^57^

The molecular hallmark of STAT1 GOF has been designated as elevated pSTAT1 (Tyr701) following type I or II IFN stimulation.^14^ However, the uniformity of this finding across all patients is perplexing in the context of high clinical variability. This suggests that elevated pSTAT1 does not fully account for STAT1 GOF disease pathogenesis. The notion that pSTAT1 may be a secondary feature of STAT1 GOF, has received support in recent years. In this regard, some studies in patient samples found STAT1 itself to be elevated, suggesting that total STAT1, rather than pSTAT1, is the primary disease driver of STAT1 GOF.^58,59^ In our current study, pSTAT1 was elevated in all GOF mutants following stimulation (as described in patients). In addition, increased total STAT1 was clearly seen in one GOF mutant, with a trend toward increased total STAT1 in two others. It is possible that elevated STAT1 levels would develop across all mutants over time following repeated stimuli.

Our analysis of gene expression at baseline and following serum starvation further supports the notion of total STAT1, rather than pSTAT1, as driving the transcriptional abnormalities seen in STAT1 GOF. Our cell models showed baseline polarization in terms of ISG expression among certain mutants, particularly E235G and P329L, suggesting that transcriptional homeostasis for some *STAT1* GOF mutants is different than that of WT. These results are recapitulated under conditions of serum starvation, suggesting that this baseline polarization may occur in a manner which is independent of external cytokine stimuli, and possibly independent (or only partially dependent) of pSTAT1. While cytokine-dependent activation of pSTAT1 has been regarded as the canonical pathway of STAT1 signaling, a well-established transcriptional role exists for unphosphorylated STAT1 (U-STAT1).^60–63^ U-STAT1 mediates the constitutive baseline activation of many ISG, in a manner which may be both cytokine-dependent and independent. It does so by acting both as a homodimer, and in complex with other transcription factors such as U-STAT2, and IRF9.^7,60–64^ It is therefore possible that mutated, unphosphorylated STAT1 molecules may result in differential transcriptional activity at baseline, causing an abnormal pattern of ISG expression even in “naïve” cells prior to cytokine stimulation. Taken together, our current findings support the notion that STAT1 GOF pathogenesis may not be fully attributed to canonical, cytokine-related pSTAT1 activation. Further work would be required to understand the mechanisms leading to baseline gene expression polarization among STAT1 GOF mutants, and what, if any, is the role of U-STAT1.

Treatment of STAT1 GOF mutants in our study with either IFNγ or IFN*α*, has shown differential stimulation responses with a mixed pattern of increased, decreased, or similar fold change in ISG expression compared to WT. Differences were noted among WT and mutants, between IFNγ or IFN*α* stimulation, and also within the same genotype and stimulation group across different genes. A case in point would be T385M, which showed no baseline differences in ISG expression compared to WT, a gain of transcriptional function with respect to all genes following IFN*α* stimulation, and no difference (and even reduced fold change for one gene) after IFNγ treatment. P329L, which was marked by increased baseline ISG expression, demonstrated reduced responsiveness to both IFNγ and IFN*α* stimuli. Comparatively, E235G, which was also characterized by enhanced baseline ISG expression, showed reduced transcriptional responses to IFNγ but increased fold change following stimulation with IFN⍰. The notion of differential transcriptional response to stimuli in STAT1 GOF is supported by previous evidence from *in-vitro* work done in patient cells. Kobbe et al showed that in T-cells from patients with the F172L mutation, fold change in *GBP1* expression was elevated compared to WT following IFN*α*, but not IFNγ stimulation, while *MIG1* fold change was elevated after treatment with either IFN*α* or IFNγ, but not combined treatment with both.^65^ Meesilpavikkai et al demonstrated that in T-cells harbouring the V653I mutation, *CXCL10* and *CD274* fold change was increased compared to WT after stimulation with IL-27, but not with IFNγ.^66^ Similar findings were reported by groups studying patient PBMC assessing the expression of various other genes.^67,68^

The intricate patterns of gene expression seen in STAT1 GOF mutants may result in part from altered STAT1-DNA binding status. For instance, the 329 residue lies one amino acid away from the site of direct DNA interaction. It is therefore conceivable that P329L could result in more constitutive baseline DNA binding and transcriptional activation, but with less potential for further transcriptional enhancement following stimulation. However, another mutation affecting the DNA binding domain, T385M, showed a vastly different gene expression pattern compared with P329L, both in terms of baseline expression and in regards to IFN reactivity, suggesting that additional factors may come into play. One such factor may relate to differential activation of STAT1-dependent and independent transcription factor complexes among the various mutants, leading to differential ISG expression both at baseline and following stimulation.

One important finding of our study relates to the designation of mutations as “GOF” or “LOF”. All in all, each STAT1 GOF mutant in our study showed evidence for transcriptional GOF with respect to at least 1 of the 5 genes measured compared to WT, either by means of increased baseline expression, or increased fold change following stimulation. However, some mutants, such as K278E and D517G showed more evidence for transcriptional LOF rather than GOF. The LOF mutant, Y701C, predictably showed reduced transcriptional responses to both stimuli across 4 of the 5 genes measured, with the exception of *OAS1* which is known to be co-regulated by non-STAT1 dependent transcriptional complexes.^6^ Our findings in STAT1 GOF mutants suggest that the molecular designation of GOF (as it relates to tyrosine phosphorylation), may not always indicate heightened gene expression. The notion of diminished ISG transcriptional responses in some mutants is supported by recent studies reporting transcriptional LOF in STAT1 GOF. Ovadia et al reported that compared with WT, the H629Y mutation showed either diminished or unchanged fold increase in gene expression in response to IFNγ.^29^ More recently, work done in a mouse model of the R274Q mutation demonstrated a reduction in the expression of the ISG *Cxcl10* and *Irf1* following viral infection *in-vivo.*^69^

The current study is constrained by a few limitations. These include a relatively small number of genes tested, and the measurement of fold change following single stimulation of naïve cells. Future work will involve larger-scale gene studies, including pathways which extend beyond the immediate group of IFN-response genes. In regards to stimulation of naïve cells, previous work showed that STAT1 GOF cells had an impaired transcriptional response not only upon initial stimulation, but also to re-stimulation.^45^ It would therefore be important in the future to study how the transcriptional responses change over time and following repeated or different stimuli. Such work may further help understand the evolution of transcriptional responses as they occur *in-vivo*. Finally, further work may be merited with regard to mechanisms resulting in differential gene expression across mutants and stimuli, and in particular elucidating the possible contribution of total or U-STAT1 to the abnormal gene expression patterns in STAT1 GOF.

In conclusion, we present a series of heterozygous *STAT1* cell models generated using CRISPR/Cas9 base-editing, showing the utility of this technique in modelling heterozygous immune-mediated disease. Our cell models demonstrate intricate patterns of ISG expression, involving transcriptional abnormalities at baseline, following serum starvation, and after stimulation with type I or II IFN. Taken together, our findings are in line with a growing body of literature suggestive of complex and multi-factorial transcriptional responses in STAT1 GOF, which cannot be simply and generally classified as either gain or loss of function. Moreover, our findings may indicate an important role for total and U-STAT1 in disease pathogenesis, in addition to the role of elevated pSTAT1. Continued investigation of gene expression patterns associated with *STAT1* mutations, may enhance our understanding of both disease pathophysiology and genotype-phenotype correlation in STAT1 GOF. This, in turn, has the potential to improve our prognostic capacity of patients affected by this disorder, and may ultimately open up new avenues for disease interrogation and targeting.

## MATERIALS AND METHODS

### Mutation selection

Previously published mutations were selected for modelling according to the following criteria: (1) transition mutations (A•G or C•T) for generation by CRISPR/Cas9 base-editing; (2) patients carrying the mutations met clinical criteria for STAT1 GOF diagnosis; and (3) previously published *in-vitro* analysis confirmed the presence of elevated Tyr701 phosphorylated-STAT1 (pSTAT1) following IFN stimulation. An additional heterozygous loss of function transition mutation, Y701C, was chosen for modelling as well.

### Cell culture

HAP1 cells (kind gift of Dr. Aleixo Muise, Toronto) were cultured in IMDM medium (Wisent Bioproducts 319-105-CL) supplemented with 10% heat-inactivated FBS (Wisent Bioproducts 080-150) and 1% penicillin-streptomycin (Wisent Bioproducts 450-201-EL). For serum-starvation experiments, HAP1 cells were placed in IMDM containing 0.25% FBS and 1% penicillin-streptomycin. Fibroblasts (ATCC PCS-201-012) for DNA-content analysis were cultured in DMEM medium (Wisent Bioproducts 319-005-CL) supplemented with 10% heat-inactivated FBS and 1% penicillin-streptomycin. Cell line authentication was done by means of Short Tandem Repeat (STR), performed by The Hospital for Sick Children The Centre for Applied Genomics (TCAG). Cells were confirmed to be Mycoplasma-free using a PCR Mycoplasma Detection Kit (Applied Biological Materials Inc. G238). Cells were cultured in a 37°C humidified incubator containing 5% CO_2._

### DNA-content analysis

DNA content analysis was performed as previously described,^70^ comparing HAP1 cells and diploid fibroblasts. Briefly, cells were washed with 1xPBS (Wisent Bioproducts 311-010-CL), trypsinized, and resuspended in complete media (1×10^6^ cells/μl), to which a low-toxicity, cell-permeable DNA dye (Vybrant DyeCycle Violet™; 1:1000, Thermo Fisher Scientific V35003) was added. Cells were incubated at 37°C for 30 minutes. Live/dead cell stain was concurrently performed using propidium iodide (1μg/μL; Thermo Fisher Scientific P1304MP), added according to manufacturer recommendation. Sample were run on BD LSR-II™ with the BD FACSDiva™ software v9.0, using the services of the Hospital for Sick Children Flow Cytometry Facility. Data were analysed using FlowJo version 10.7.1. Median Fluorescence Intensity (MFI) peaks were compared for HAP1 and fibroblasts at G_0_/G_1_ and G_2/_M.

### Cloning

The sgRNA vector, BPK1520_puroR, a plasmid containing a cloning site for sgRNA under control of a U6 promoter, as well as a puromycin resistance cassette, was generated as previously described.^35^ The following oligonucleotides were used to clone the sgRNA required for mutant generation: E235G, Fwd CACCGATGAACTAGTGGAGTGGAAG, Rev AAACCTTCCACTCCACTAGTTCATC; K278E, Fwd CACCGCTTAAAAAGTTGGAGGAAT, Rev AAACATTCCTCCAACTTTTTAAGC; P329L, Fwd CACCGCTGAGGGTGCGTTGGCATGC, Rev: AAACGCATGCCAACGCACCCTCAGC; T385M: Fwd: CACCGGGCACGCACACAAAAGTGA, Rev: AAACTCACTTTTGTGTGCGTGCCC; D517G, Fwd: CACCGTGTGGACCAGCTGAACATGT, Rev: AAACACATGTTCAGCTGGTCCACAC; Y701C, Fwd: CACCGTGGATATATCAAGACTGAGT, Rev: AAACACTCAGTCTTGATATATCCAC.

Oligonucleotides were annealed using an annealing buffer (10 mM Tris, pH 7.5 - 8.0, 50 mM NaCl, 1 mM EDTA) and phosphorylated using T4 Polynucleotide Kinase (New England BioLabs M0201L) according to manufacturer recommendations. BPK1520_puroR was linearized with BsmBI (New England BioLabs R0580S) and dephosphorylated using recombinant shrimp alkaline phosphatase (New England BioLabs M0371L). Annealed and phosphorylated oligonucleotides were cloned into linearized and dephosphorylated BPK1520_puroR using T4 DNA Ligase (New England BioLabs M0202L) according to manufacturer recommendations. Subsequently, One Shot™ TOP10 Chemically Competent E. coli (Thermo Fisher Scientific C404003) were transformed with the ligation products and plated on LB-Ampicillin agar plates (50 μg/mL ampicillin). Resultant colonies were inoculated overnight in LB-ampicillin, and plasmids purified using the QIAprep Spin Miniprep Kit (Qiagen 27106) according to manufacturer recommendations. The following plasmids were used for the purpose base-editing: pCMV_ABEmax_P2A_GFP (gift from Dr. David Liu, Addgene plasmid 112101) was used to generate E235G, K278E, P329L, and D517G. Plasmids pCAG-CBE4max-SpG-P2A-EGFP and pCMV-T7-ABEmax(7.10)-SpRY-P2A-EGFP (gifts from Dr. Benjamin Kleinstiver, Addgene plasmids 139998 and 140003) were used to generate T385M and Y701C, respectively.

### Transfection and selection

Twenty-four hours prior to transfection, 4×10^5^ cells were seeded in a 6-well plate. The following day, cells were transfected with 2500ng of total DNA, containing the Cas9 base-editor expression vector and the sgRNA expression vector containing the PuroR gene, at a 1:1 ratio (w/w). Transfection was performed using Lipofecatmine™ 3000 Transfection Reagent (Thermo Fisher Scientific L3000001) according to manufacturer recommendations. To enrich for transfected cells, 24 hours post-transfection cells were subjected to puromycin selection (0.7 μg/mL; Thermo Fisher Scientific A1113803) for 72 hours. Following puromycin selection, estimation of base-editing efficiency in the bulk population was performed as follows: ~1×10^6^ cells were collected from which genomic DNA was isolated using the DNeasy Blood & Tissue Kit (Qiagen 69506). This was followed by PCR amplification of the desired region using DreamTaq Polymerase (Thermo Fisher Scientific EP0705). Amplified DNA was PCR-purified using QIAquick PCR Purification Kit (Qiagen 28106) following the manufacturer’s protocol and prepared for Sanger sequencing using the BigDye™ Terminator v3.1 Cycle Sequencing Kit (Thermo Fisher Scientific 4337457). Samples were sequenced on Applied Biosystems SeqStudio Genetic Analyzer (Thermo Fisher Scientific), and sequencing AB1 files were input into the online base-editing analysis tool, editR.^71^ This provided an estimated percentage editing of the bulk population, as well as percentage editing (if any) of any adjacent bases.

### Single-cell sorting and clone screening

Following estimation of editing efficiency, cells were trypsinized and resuspended in FACS buffer (1xPBS without calcium and magnesium pH 7.4, supplemented 2% FBS and 2.5mM EDTA) at a concentration of 1×10^6^ cells/mL. Live/dead cell stain was performed using propidium iodide. Live single cells were sorted on MoFloXDP Cell Sorter (Beckman Coulter), using the services of the Hospital for Sick Children Flow Cytometry Facility. Cells were sorted into a 96-well plate containing full media and allowed to clonally expand for a period of 14 days. Following a 2-week recovery period, single-cell clones underwent genomic DNA isolation, followed by PCR amplification, purification and Sanger sequencing as described above.

### Immunoblotting

Immunoblotting was used to determine STAT1 and pSTAT1 protein levels in IFNγ-stimulated or unstimulated cells, over 5 independent experiments. For each experiment, 4×10^5^ cells were seeded in a 6-well plate. Twenty-four hours later, Human Recombinant IFNγ (10ng/mL, StemCell Technologies 78020) was added to the media for a period of 60 minutes. Cells were subsequently washed twice with cold 1xPBS, and whole-cell lysates were obtained by lysing cells in RIPA Lysis and Extraction Buffer (Thermo Fisher Scientific 89900), supplemented with Halt™ Protease and Phosphatase Inhibitor Cocktail (Thermo Fisher Scientific 78440), on ice for 30 minutes. Lysates were sonicated and then centrifuged at 12,000xg for a period of 15 minutes at 4LJC. Protein concentrations were determined using the Pierce™ BCA Protein Assay Kit (Thermo Fisher Scientific 23552). Samples were then prepared by addition of Nu PAGE™ LDS Sample Buffer (4X) (Thermo Fisher Scientific NP0007) followed by boiling at 100LJC for 5 minutes. Samples were subjected to SDS-Page separation by running 20μg of total protein on a NuPage 4-12% Bis-Tris gel (Thermo Fisher Scientific NP0336BOX) using NuPAGE™ MOPS SDS Running Buffer (Thermo Fisher Scientific NP000102). For each experiment, samples for pSTAT1 and STAT1 were run in parallel. Protein was subsequently transferred to a nitrocellulose membrane using the iBlot 2 Dry Blotting System (Thermo Fisher Scientific). Following transfer, membranes were blocked in 1xTris-Buffered Saline (50 mM Tris-Cl, pH 7.5. 150 mM NaCl) containing 5% bovine serum albumin (Sigma Aldrich A7906-50G) for 1 hour at room temperature. Membranes were then incubated at 4◻C overnight with primary antibodies against pSTAT1 (pY701; clone D4A7; Cell Signaling 7649S), total STAT1 (Clone D1K9Y, Cell Signaling 14994) or alpha-tubulin (Clone DM1A; Sigma Aldrich T6199-100UL). The following day, membranes were washed in tris-buffered saline and incubated for 1 hour at room temperature with one of the following secondary antibodies: Donkey anti-Rabbit IgG (H+L) Highly Cross-Adsorbed Secondary Antibody, Alexa Fluor 647 (Thermo Fisher Scientific A-31573) or Donkey anti-Mouse IgG (H+L) Highly Cross-Adsorbed Secondary Antibody, Alexa Fluor 647 (Thermo Fisher Scientific A-31571). Membranes were imaged using ChemiDoc MP imaging system (Bio-Rad) and analyzed with Image Lab software (^©^2017 Bio-Rad Laboratories; version 6.0.1).

### RNA isolation and Quantitative real-time PCR

RNA analysis was performed to determine gene expression levels across the different genotypes under various conditions. Experiments were repeated a minimum of 5 times for each gene, in at least technical duplicates, and pooled data for each gene was collected. To determine baseline gene expression levels, cells were grown in full media and harvested without stimulation once reaching 70-80% confluence. For determination of baseline gene expression under low-serum conditions, cells were seeded as described above and allowed to adhere in full media for a period of 24 hours. Cells were subsequently thoroughly washed with PBS and placed in low-serum media (IMDM+0.25% FBS) for an additional 24 hours. For stimulation experiments, cells in full media were incubated with Human Recombinant IFN*α*-2A (10ng/mL, StemCell Technologies 78076.1) or IFNγ (10ng/mL) for 6 hours prior to harvesting and RNA extraction using RNeasy Mini Kit (Qiagen 74106). Next, 1000ng of RNA was reverse-transcribed using SuperScript™ III First-Strand Synthesis System (Thermo Fisher Scientific 18080051) following the manufacturer’s protocol. Quantitative real-time PCR (qRT-PCR) using PowerUp™ SYBR™ Green Master Mix (Thermo Fisher Scientific A25742) was performed on an Applied Biosystems QuantStudio 3 Real-Time PCR System (Applied Biosystems). Quantification of the following genes was done: *GBP1, IFIT2, IRF1, APOL6, OAS1*, with *GAPDH* used as housekeeping control. The following primers were used, designed using Primer3 (v.0.4.0): *GBP1* Fwd AGGAGTTAGCGGCCCAGCTAGAAA, Rev AAAATGACCTGAAGTAAAGCTGAGC; *IFIT2* Fwd GCACTGCAACCATGAGTGAGA, Rev CAAGTTCCAGGTGAAATGGCA; *IRF1* Fwd TCCTGCAGCAGAGCCAACATGCCCA, Rev CCGGGATTTGGTTGGAATTAATCTG; *APOL6* Fwd TTGGTTTGCAAAGGGATGAGGATGA, Rev TCTTTCAATCTGGGAAATTCTCTCA; *OAS1* Fwd CAAGGTGGTAAAGGGTGGCTCCTCA, Rev TAACTGATCCTGAAAAGTGGTGAGA; *GAPDH* Fwd CAATGACCCCTTCATTGACCTC, Rev GATCTCGCTCCTGGAAGATG. The relative expression levels were compared using the ΔΔCt method.

### Statistical analysis

Graphical data were represented as means + standard error of mean. Statistical analysis was performed using GraphPad Prism 8 (version 8.4.3). One-way ANOVA with Dunnett’s post-hoc test was used to determine differences among mutants and wildtype. Statistical significance was represented as: *p < 0.05, **p < 0.01, ***p < 0.001, ****p < 0.0001.

## Data availability

The data that support the findings of this study are available from the corresponding author upon reasonable request.

## Acknowledgments

Funding this work was provided by Immunodeficiency Canada (OS). Salary support for OS has been provided by the Ontario Ministry of Health Clinician Investigator Program, the Hospital for Sick Children Clinician Scientist Training Program, and the Canadian Child Health Clinician Scientist Program. Figures 1 and 3 were created with Biorender.com.

## Author Contributions

Study conception and design were done by OS, CMR, EAI, RDC. Experimental data acquisition was performed by OS, KL. Data analysis was performed by OS, KL, SE. Data interpretation was done by OS, SE, CMR, EAI, RDC. Writing of first draft was performed by OS. Further writing, reviewing and editing was by OS, KL, SE, CMR, EAI, RDC. The work was jointly supervised by EAI and RDC. All authors have approved the submitted version and have agreed both to be personally accountable for their own contributions and to ensure that questions related to the accuracy or integrity of any part of the work, even ones in which the author was not personally involved, are appropriately investigated, resolved, and the resolution documented in the literature.

