## Supplementary Figure for "Heterozygous Cell Models of STAT1 Gain-of-Function Reveal a Broad Spectrum of Interferon-Signature Gene Transcriptional Responses"

**Supplementary Figure Legends**

**Figure 1.** DNA content analysis of demonstrates diploidy of HAP1 cells. | **(A)** DNA content was measured in HAP1 (blue), wildtype human fibroblasts (known to be diploid; orange) and a 1:1 mixture of fibroblasts and HAP1 cells (red). DNA content was assessed by measuring the median fluorescence intensity (MFI) of a live cell DNA dye (Vybrant DyeCycle Violet^TM^) across the cell cycle. MFI ratios between HAP1 and fibroblasts were calculated for the G_0_/G_1_ phase (left peak) and G_2_/M phase (right peak). An MFI ratio of ~1 for both peaks indicate that the ploidy of HAP1 cells is identical to that of diploid fibroblasts. **(B)** Gating strategy used for DNA content analysis. Following exclusion of debris in the FSC-A/SSC-A window, doublets were excluded using FSC-A/FSC-H followed by the SSC-A/SSC-H parameters. Dead cells were excluded by plotting the G660-A (propidium iodide) fluorescence intensity vs. SSC-H. Events associated with G660-A fluorescence intensity higher than 10^3 were excluded. Finally, for all live single cells, fluorescence intensity ofV450-A (Vybrant DyeCycle Violet^TM^) was plotted as a histogram.

**Figure 2.** Uncropped immunoblot membranes presented in main manuscript. | (A) uncropped immunoblot membrane presenting expression of pSTAT1 (Tyr701) with alpha-tubulin loading control, following 60 minutes of stimulation with IFNγ (10ng/mL); (B) uncropped immunoblot membrane presenting total STAT1 with alpha-tubulin loading control following 60 minutes of stimulation with IFNγ (10ng/mL).

**Supplementary Figures**

**Supplementary Figure 1:**

**A**

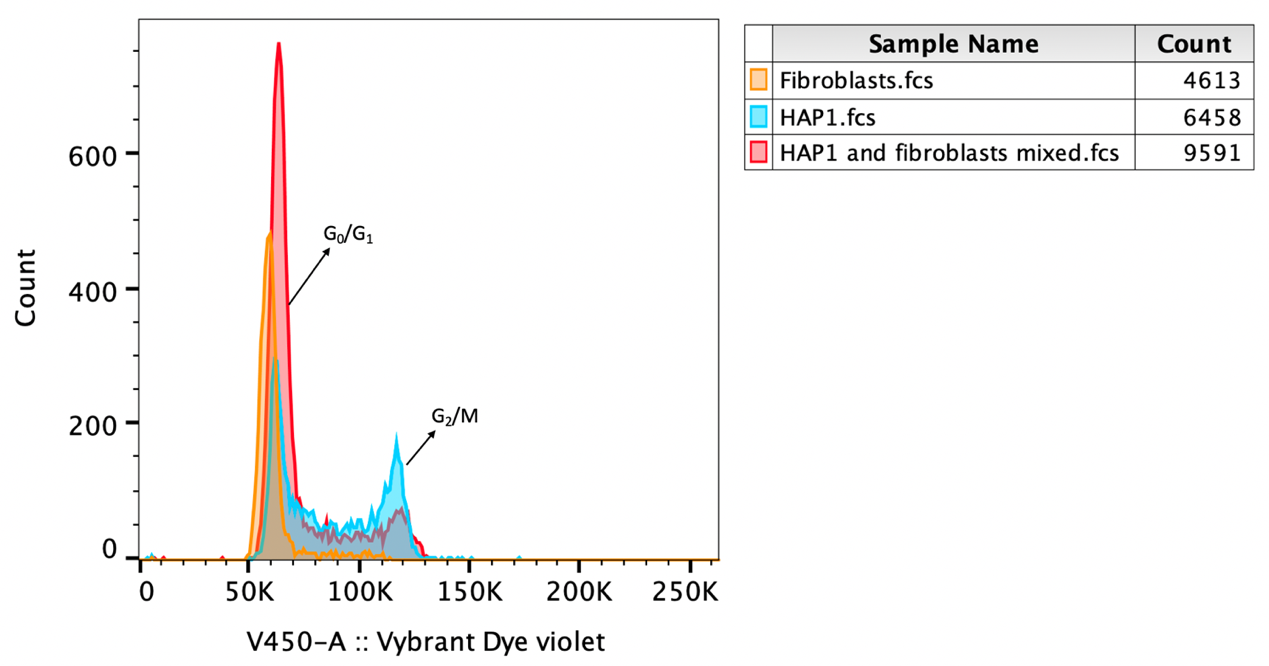

**B**

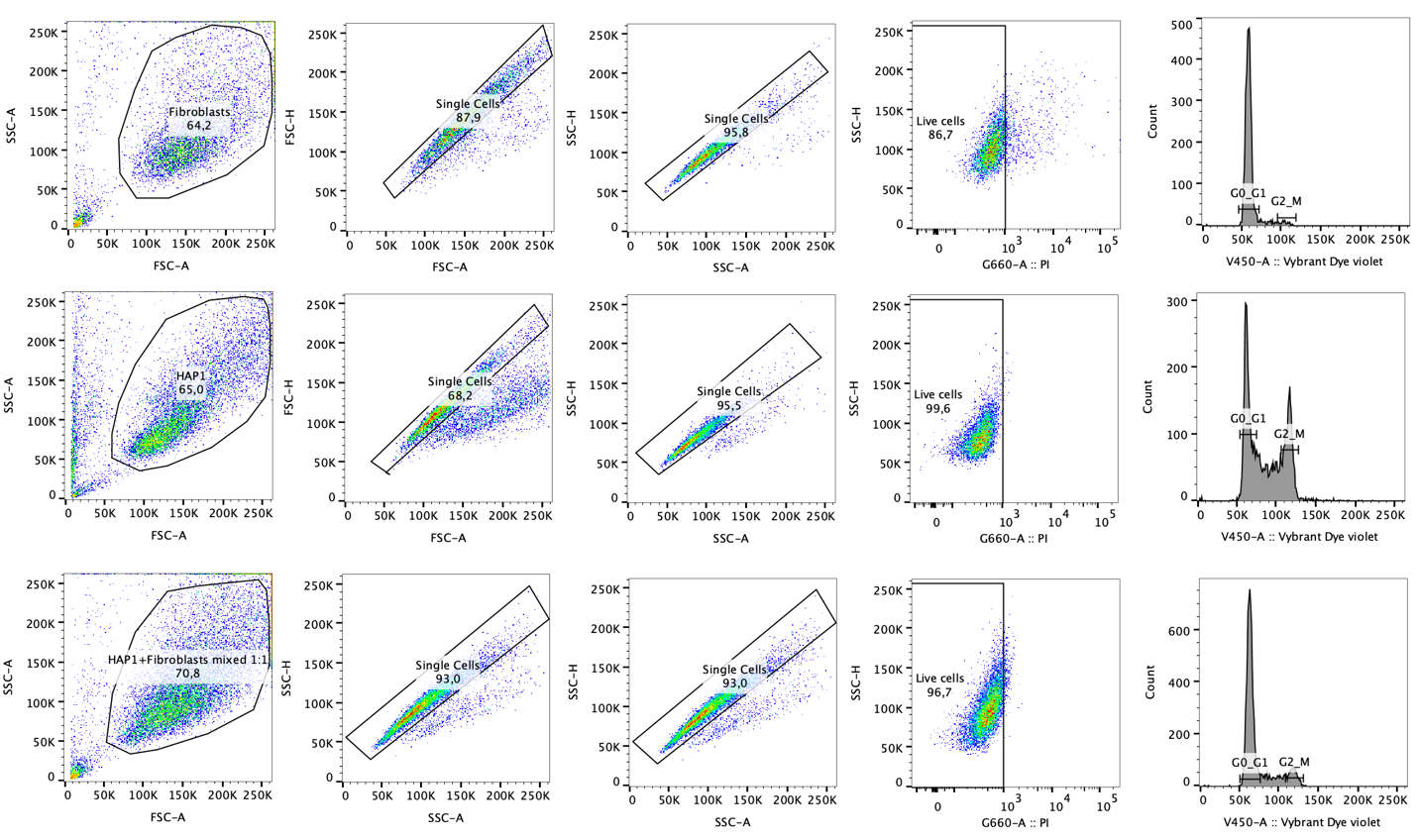
**Supplementary Figure 2:**

**A**

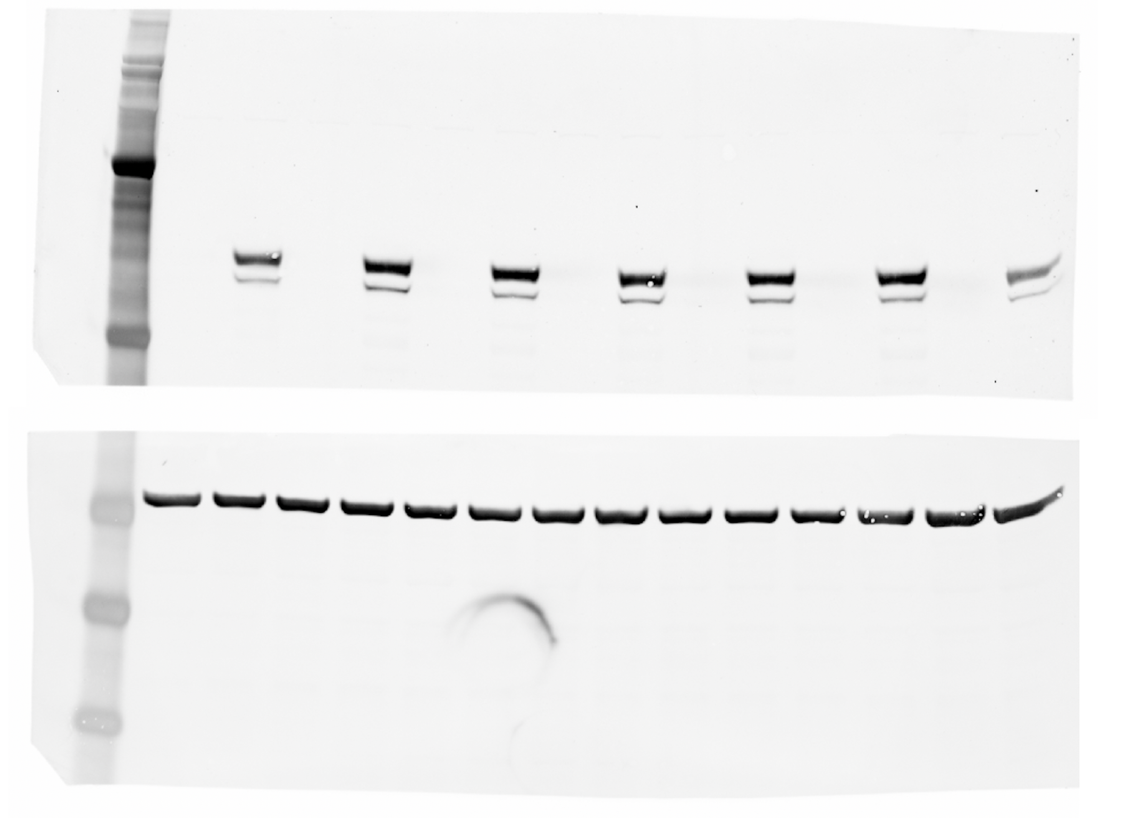

**B**

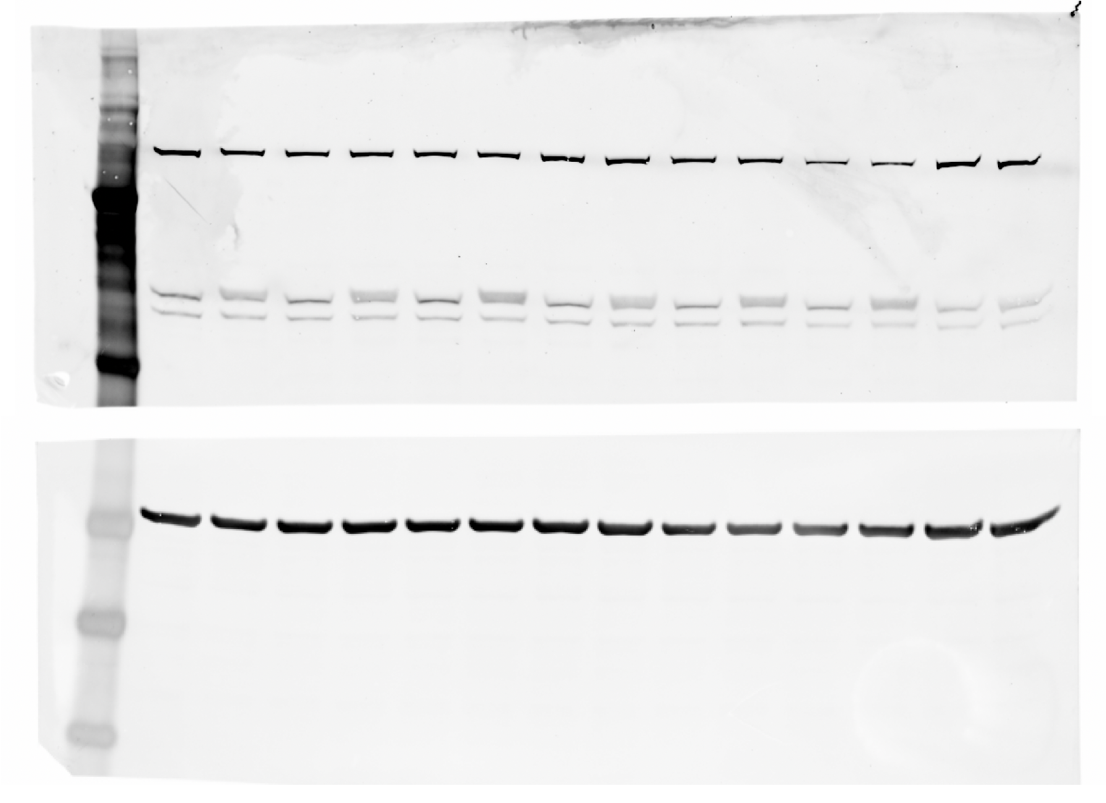
